# The demographic buffering strategy has a threshold of effectiveness to increases in environmental stochasticity

**DOI:** 10.1101/2020.05.15.098095

**Authors:** R.C. Rodríguez-Caro, P. Capdevila, E. Graciá, J. M. Barbosa, A. Giménez, R. Salguero-Gómez

## Abstract

1. Animal populations have developed multiple strategies to deal with environmental change. Among them, the demographic buffering strategy consists on constraining the temporal variation of the vital rate(s) (e.g., survival, growth, reproduction) that most affect(s) the overall performance of the population. Given the increase in environmental stochasticity of the current global change scenario, identifying the thresholds beyond which populations are not able to remain viable -despite their potential buffering strategies- is of utmost importance.
2. Tortoises are known to buffer the temporal variation in survival (*i.e.* this vital rate has the highest contribution to the population growth rate *λ*) at the expense of a high variability on reproductive rates (lowest contribution to *λ*). To identify the potential threshold in buffering ability, here we use field data collected across a decade on 15 locations of *Testudo graeca* along South-Eastern Spain. We analyse the effects of environmental variables (precipitation, temperature, and NDVI) on the probability of laying eggs and the number of eggs per clutch. Finally, we couple the demographic and environmental data to parametrise integral projection models (IPMs) to simulate the effects of different scenarios of drought recurrence on population growth rate.
3. We find that droughts negatively affect the probability of laying eggs, but the overall effects on the population growth rates of *T. graeca* under the current drought frequencies (one per decade) are negligible. However, increasing the annual frequency of droughts decreases the buffering ability of *T. graeca* populations, with a threshold at three droughts per decade.
4. Although some species may buffer current environmental regimes by carefully orchestrating how their vital rates vary through time, a demographic buffering strategy may alone not warrant population viability in extreme regimes. Our findings support the hypothesis that the buffering strategy indeed has a threshold of effectiveness. Our methodological approach also provides a useful pipeline for ecologists and managers to determine how effective the management of environmental drivers can be for demographically buffering populations, and which scenarios may not provide long-term species persistence.

## Introduction

Persistence under changing environmental conditions is one of the main challenges faced by most species worldwide (Tuljapurkar, 1990). Environmental stochasticity is predicted to increase with global climate change (Hansen et al., 2006; Alexander et al., 2006; IPCC 2014). This environmental stochasticity can cause the decline of natural populations and lead to their extinction due to the physiological and demographic responses of organisms to unexpected climate conditions (Chown, et al., 2010; Selwood, McGeoch & Mac Nally, 2014; Lawson, Vindenes, Bailey, & van de Pol, 2015). Thus, understanding how extant life history strategies may respond to future changes, such as projected increases of droughts (Burke, Brown & Christidis, 2006) is critical (Coumou & Rahmstorf, 2012; McDonald et al., 2017). This knowledge is particularly needed -yet challenging- in threatened species because of their frequently small population sizes and constrained distributions (Parmesan & Yohe, 2003, Thomas et al., 2004). Demographic descriptors, such as population size (*N*), ultimately control the degree of environmental stochasticity that a given population can cope with before going locally extinct (Lanfear, Kokko & Eyre-Walker, 2014). For instance, reductions in *N* via negative effects of environmental stochasticity on underlying vital rates (Tuljapurkar, 1990) can lead to the population collapse if *N* reaches values lower than the effective population size *N_e_* of the species (Shaffer, 1981).

Species have evolved a wide range of life history strategies to allow them to persist in different environments (Pianka, 1970; Southwood, 1988; Stearns, 1992; Lytle, 2001; Morris & Doak, 2004; Boyce et al., 2006). Examples of such strategies include the large offspring size driven via parental care, which enhances establishment (Shine, 1978); or self-pollination in low-density populations, which increases short-term demographic viability (Kalisz, Vogler, & Hanley, 2004). This diversity of strategies ultimately emerges from the limitations imposed by finite resources and physiological constraints (Stearns, 1992). During the last decades, researchers have focused on classifying such strategies according to investments on specific vital rates (e.g. fast-slow or reproductive strategies; Gaillard et al., 1989; Stearns, 1992; Bielby et al., 2007; Salguero-Gómez, 2017; Healy, Ezard, Jones, Salguero-Gómez & Buckley, 2019). However, these classifications do not explicitly consider how the investments in each vital rate may vary with time and the implications for population viability (but see Pfister 1998; Doak & Morris, 2010).

Evidence is starting to emerge regarding a continuum of strategies to deal with environmental variation that ranks species from highly demographically buffered to highly demographically labile (Gillespie, 1977; Pfister, 1998; Koons, Pavard, Baudisch & Metcalf, 2009). Demographically labile species, such as some annual plants (Levine and Rees, 2004), persist in stochastic environments by allowing the most sensitive vital rate(s) (i.e. the vital rate(s) with large effect on population growth) to vary with environmental conditions, thus “tracking the environment” (Koons et al., 2009; Jongejans, De Kroon, Tuljapurkar & Shea, 2010). In contrast, demographically buffered species, such as primates (Campos et al., 2017), tend to constrain the temporal variation of the vital rates that most affect population growth rate *λ* (Pfister, 1998; Boyce et al., 2006). A next logical step in this research agenda is to identify thresholds beyond which demographic strategies may no longer be able to cope with increasing environmental stochasticity (Doak and Morris, 2010).

Chelonians are known to demographically buffer changes in environmental conditions (Heppell, 1998). This taxonomic group includes some of the longest-lived tetrapods, such as Testudinidae or Emydidae, with lifespans of over 100 years (Castanet, 1994). Chelonians are ideal organisms to investigate the possibility of thresholds of the buffering strategy under harsh climate conditions because they are ectothermic and therefore their vital rates are strongly influenced by the environment (Ihlow et al., 2012). Terrestrial tortoises exhibit high temporal variability in their reproduction output and has a low contribution to population growth, *λ* (Congdon, 1989; Doak, Kareiva & Klepetka, 1994; Wisdom, Mills & Doak, 2000). Reproductive rate in chelonians is also usually diminished due to adverse environmental changes, such as reductions in rainfall regimens or increases in temperature (Turner, Hayden, Burge & Roberson, 1986; Nieuwolt-Dacanay, 1997). However, tortoise populations buffer such negative impacts on reproductive rates through a constant, high adult survival rates through time (e.g. Turner et al., 1986; Henen, 1997). Still, the projected increases in the frequency of adverse environmental conditions (Seneviratne et al., 2012) poses novel challenges for the viability of tortoise populations. Thus, identifying the limits of their buffering ability is crucial for their effective conservation, as well as for that of any demographically buffering species.

Here, we evaluate the population viability of a chelonian species, the spur-thighed tortoise (*Testudo gracea*), in SE Spain (Sanz-Aguilar et al., 2011; Rodríguez-Caro, Graciá, Anadón & Giménez., 2013; Rodríguez-Caro et al., 2019; Graciá et al., 2020). We hypothesised that the projected increase in adverse condition frequency in the region will push the species beyond its buffering ability and trigger local extinctions. Using field demographic data collected along a decade and containing radiographed dorso-ventral body area together with bibliographic information on this species, we parameterise an integral projection model (IPM; Easterling, Ellner, & Dixon, 2000) to: (1) identify the effects of environmental conditions in reproduction rates; (2) evaluate the effects of current and projected adverse conditions (*i.e.* droughts) on its population growth rates (*λ*) to determine the limit of its demographic buffering; and (3) identify the demographic process(es) that is/are most responsible for a decrease in population performance due to increases in adverse conditions (*i.e.* droughts) using perturbation analyses.

## Methods

### Study system

The spur-thighed tortoise, *Testudo graeca* L. (Testudinidae) is a long-lived endangered species (*Vulnerable*, IUCN, 2012). This species is distributed across North Africa, Southern Europe, and Southwest Asia (Graciá et al., 2017a). The largest population of *T. graeca* in Spain are located in the South-East of the Iberian Peninsula (Fig. 1a). In this area, *T. graeca* are distributed across ~2,600 km^2^ of semiarid mountains, where annual precipitation ranges between 150 and 570 mm/yr. This species has slow population dynamics (*sensu* Wilbur & Morin, 1988), steady population growth rates (*i.e.*, *λ* ~ 1; Rodríguez◻Caro, Lima, Anadón, Graciá, & Giménez, 2016), delayed maturation (9-12 years; Graciá et al., 2020), and low offspring production (< 2 hatching per female per year; Jiménez-Franco et al., 2020). The life cycle of the species includes three stages: juveniles, subadults, and adults (Fig 1b). The survival of juveniles is typically low due to their soft carapace coupled with high predation by wild boars, rats or foxes (Lambert, 1982; Hailey, 1988, Díaz-Paniagua, Keller & Andreu, 1997), whereas in subadults (4-8 years old) and adults (reproductive individuals) survival is much higher (0.79-0.98; Sanz-Aguilar et al., 2011, Rodríguez-Caro et al., 2013).

**Fig 1.**
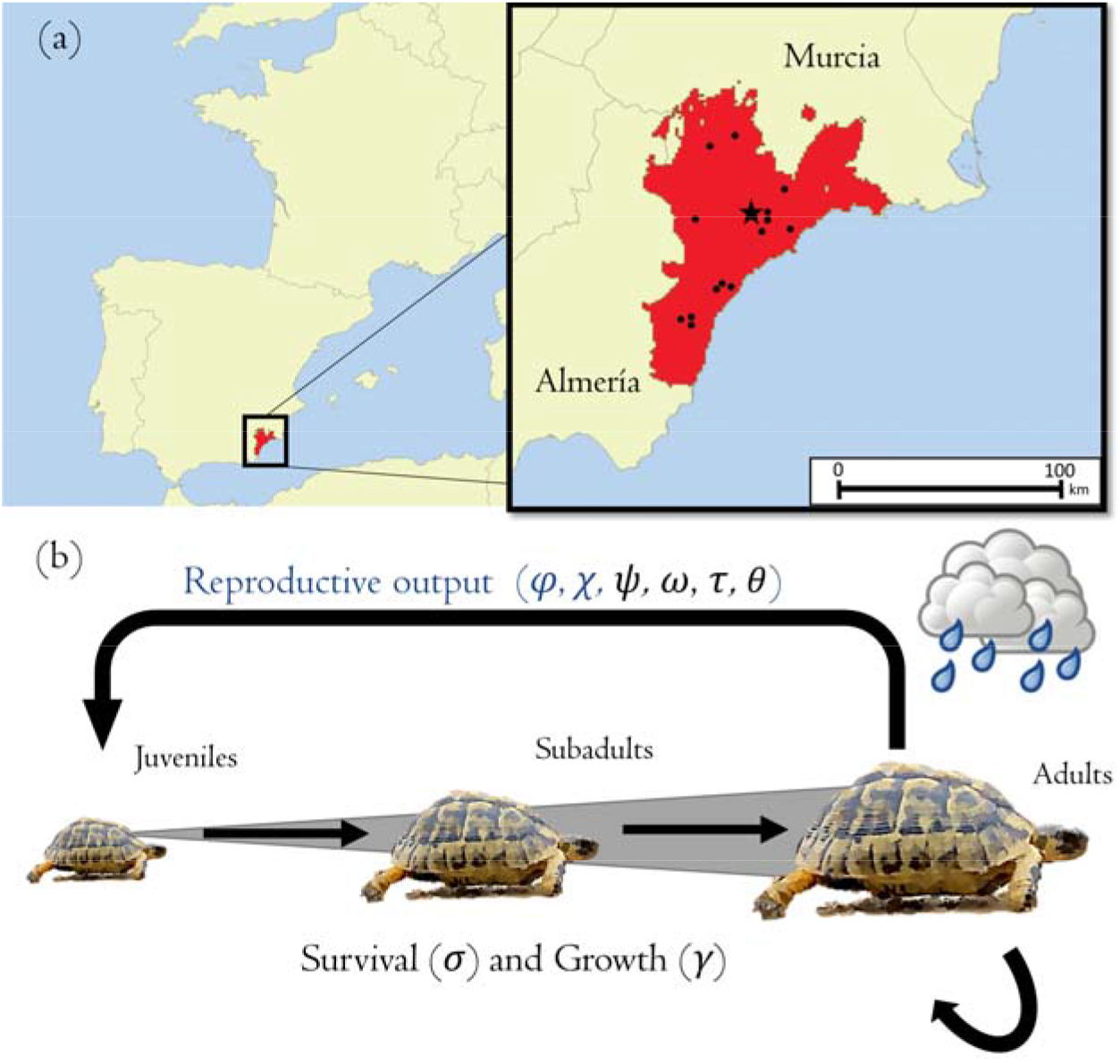
(a) Distribution of the spur-thighed tortoise (*Testudo graeca*) in the south-eastern Iberian Peninsula (red). Black dots represent the locations of sampled sites, while the black star represents the location of the study site Galera (Sierra de la Carrasquilla, Murcia, Spain 37°32’N, 1°39’W), for which we developed the drought simulations. (b) Life cycle of T. *graeca*, including three stages (juveniles, subadults, and adults), and the different demographic processes used in our Integral Projection Models. Individuals may grow to a given size with a probability *γ* if they survive (*σ*) from one year to the next. Only adults can reproduce (blue arrow). To quantify reproduction, we estimated the probability of reproduction (*φ*), the number of eggs produced per adult per clutch (*χ*), the number of clutches per year (*ψ*), the probability of hatching (*ω*), the probability of surviving to the one year (*τ*), and the size distribution of offspring the next year (*θ*) from field data. The vital rate parameters shown in blue are known to be affected by environmental conditions, and thus they are the focus of our study.

To obtain reproductive information for this species, we conducted fieldwork during a decade (specifically, in 2006-2007, 2010-2011, and 2014-2016), across 15 sites (Fig 1a), sampling a total of 634 female individuals (Appendix 1). The timing of the sampling coincided with the peak of the reproductive season, in April-June, so that adult female individuals could be assessed for their reproductive status (Díaz-Paniagua, Keller & Andreu, 1996). We captured individuals manually along line transects of 2 km, repeated three times per site, during the spring (Rodríguez-Caro et al. 2017). For each individual, we measured its carapace length (CL) and femoral width (FW) with digital callipers. Adults were sexed according to their morphological characteristics – juveniles and subadults do not show sexual dimorphism (López-Jurado, Talavera-Torralba, Ibánez-González, MacIvor & García-Alcaza, 1979). Females and subadults are morphologically indifferentiable other than by size, and so we identified the minimum size in femoral width (FW_m_) above which females lay eggs. All females were radiographed dorsoventrally with a portable X-ray machine set at 60kV (20mAs) at a distance of 1 m (Hinton, Fledderman, Lovich, Congdon, & Gibbons, 1997), following methods described in Gibbons and Green (1979). In the radiography, we identified whether females carried eggs to estimate the probability of reproduction (*φ*) and, if so, counted the number of eggs (*χ*; Fig. 1b). We considered only females larger than FW_m_ = 61 mm for further analysis (N = 534 females) because this was the minimum observed size in our radiographic examinations of a female with eggs.

To parameterise our demographic model, we used survival (*σ*) and growth (*γ*) estimates from previous studies at the Galera site (Sierra de la Carrasquilla, Murcia, Spain 37°32’N, 1°39’W; Fig. 1). Briefly, at that location, Sanz-Aguilar et al. (2011) estimated survival rates using 11 years of capture-recapture data (1999-2009) and 1,009 observations across 675 individuals of the same species. For our study, we used the survival rates estimated in 1999-2004 by Sanz-Aguilar et al. (2011): 0.20 (0.08–0.42, 95% C.I.) for juveniles, 0.79 (0.57–0.90) for subadults, and 0.98 (0.92–0.99) for adults, because this period was not affected by disturbances. Rodríguez-Caro et al. (2013) studied body growth patterns at the same site in a similar period (2000-2004). In it, 36 individuals were measured between captures-recaptures, and growth rates were modelled using von Bertalanffy regression model. We used this information to quantify individual growth rates based on carapace length (CL).

### Environmental variables: temperature, precipitation, and vegetation greenness

To examine the relationship between environmental variables and reproductive rates, we calculated the average climatic and vegetation productivity at each of the 15 sites (Fig. 1a). We selected a spatial scale of 500 m radio to represent (i) local conditions at each site, (ii) regional conditions aiming to account for the landscape heterogeneity in the surroundings of the study area, and (iii) the habitat characteristics without diluting nearby areas with different land uses (Anadón et al., 2006). For each site, we calculated mean annual temperature (°C), total annual rainfall (mm), and the annual mean of Normalised Difference Vegetation Index (NDVI), an indicator of the primary productivity and biomass (Rouse, Haas, Schell & Deerin, 1974) as food resource for herbivores (Pettorelli et al., 2011). These variables were obtained annually between 1995 and 2017. Temperature data, with a 5 km spatial grid resolution, were obtained from the Oxford Daytime Land Surface Temperature dataset (Weiss et al., 2014). This temperature product is based on the Moderate Resolution Imaging Spectroradiometer (MODIS) land surface temperature data (MOD11A2), which was gap-filled to eliminate missing data caused by factors such as cloud cover using methods described in Weiss et al. (2014). Precipitation data were obtained from the Climate Hazards Group InfraRed Precipitation with Station data (CHIRPS; Funk et al., 2015), a quasi-global gridded rainfall time series with 0.05° spatial resolution. We calculated NDVI (spatial resolution of 250 m) using the MYD13Q1 V6 product (NASA Land Processes Distributed Active Archive Center). This MODIS NDVI product is computed from atmospherically corrected bi-directional surface reflectance that have been masked for water, clouds, heavy aerosols, and cloud shadows. Per-pixel Quality Assessment metadata (Huete et al., 2002) were used to check product performance to measure NDVI among study sites and across years.

### Statistical analysis of environmental variables

To identify the environmental drivers of population performance of *T. graeca*, we used a moving window approach (van de Pol et al., 2016). Briefly, this approach consists in testing the effect of a given environmental factor under different temporal windows on a chosen response variable and comparing these models to identify potential environmental signals (Bailey & van de Pol, 2016). We carried out this analysis using the package *climwin* (Bailey & van de Pol, 2016) in R (R Core Team 2019). We fitted each of the environmental variables (temperate, precipitation and NDVI in different time windows) to the reproduction data (probability of being reproductive, *φ*; and number of eggs, *χ*; Fig. 1a). We compared the models to identify multiple (short- and long-lag) signals within the same climatic variable. We determined a baseline model structure for reproduction without environmental effects as a null hypothesis for *φ* and for *χ*, and we then tested the importance of several time windows. The time windows were measured in months, ranging in duration from 0 (*i.e.*, same time as vital rate measurement) to 12 months (*i.e.*, a year prior to the observation). We used an absolute time window method before the hypothetical censuses on 1 June of each year and relative time windows with the real day of the census. We used generalised linear models with a binomial (“logit” link) and Poisson family (“log” link), for *φ* and *χ*, respectively. Last, we compared the *climwin* model outputs, and selected the best model as that with the lowest AIC while having ΔAIC>2, or the simplest model of a range of models with lowest AIC but with ΔAIC<2 (Burnham and Anderson, 2002).

We found a strong positive effect of precipitation on the probability of reproduction (*φ*, the best eight models are described in Appendix 2). However, we did not find a significant effect of environmental variables (precipitation, temperature, or NDVI) on clutch size (*χ*). We found that the range of action of each environmental variable (temperature, precipitation, and NDVI) on the probability of reproduction *φ* was significantly different. The positive effects of precipitation on the probability of reproduction *φ* were most conspicuous during the three months prior to the observation, while the negative effects of temperature included the full year prior to the vital rate measurement. On the contrary, the positive effect of NDVI included the last 6-7 months prior to the observation. Out of these climatic variables, spring precipitation prior to the sampling month raised a more parsimonious model (ΔAIC between −62.3 and −58.3) than temperature (ΔAIC of best model was −45.6) or NDVI (ΔAIC of best model was −33.6; Appendix 2). Based on these analyses, we decided not to build multivariable models to predict vital rates as a function of these environmental variables since these variables were highly correlated (Pearson’s *r*> 0.75, *p*<0.05).

### Demographic modelling

To determine a potential threshold of drought recurrence beyond which this species is not able to effectively buffer against environmental stochasticity, we parameterised a stochastic integral projection model (IPM; Easterling et al., 2000). An IPM describes the structure *n* and dynamics of individuals in a population classified by one or more continuous traits *z’s* during the time period *t* to *t*+1 (eqn. 1; Ellner & Rees, 2006). We parameterised the IPMs coupling the field and bibliographic demographic data, together with climatological data, as described above, to analyse the effects of projected environmental changes on population dynamics. The single continuous state variable *z* in our IPM was carapace length (CL, mm), which is continuous over the three life stages (Fig. 1) using a similar framework to Rodríguez-Caro et al. (2013). The overall IPM is described by equation 1, where the kernel *K* represents the overall dynamics of the population. The *K* kernel can be sub-divided in the subkernels *P* and *F* (equation 2).

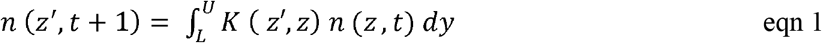

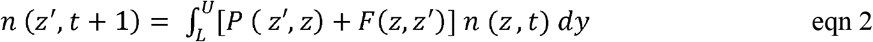

The subkernel *P* describes how individuals may change in trait *z* value (*γ*, growth), conditional on survival (*σ*) (equation 3). The subkernel *F* describes the contribution of reproductive individuals of size *z* at time *t* to new recruits of a given size *z’* at *t*+1 (equation 4). Here, the subkernel *F* is defined by the six vital rate functions described in the reproductive route of Figure 1.b: probability of reproduction in year *t* (*φ*), number of eggs per clutch (*χ*), number of clutches per individual in time *t* (*ψ*), probability of egg hatching in time *t* (*ω*), survival probability of the offspring from *t* to *t*+1 (*τ*), and size distribution of offspring in *t*+1 (*θ*). As we developed this model just for females, we halved egg production under the assumption that the female-male sex ratio is 1:1, because the sex ratio in populations of the South-East of Spain is typically 1:1 (Graciá et al, 2017b). The vital rates *φ* and *χ* were estimated empirically in this study, and the remaining four (*ψ*, *ω*, *τ*, and *θ*) were obtained from previous studies on the same species in other population of Spain (Díaz-Paniagua et al., 1997, Keller, Díaz-Paniagua and Andreu, 1997).

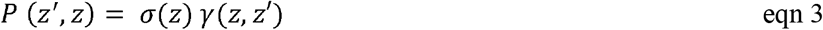

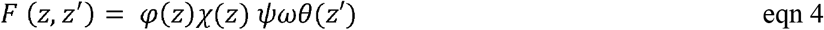

The limits of the integral of *K* in equations 1 and 2 were initially evaluated as the minimum observed size (*L* = 27 mm) to the maximum observed size (*U*=176 mm) across all sites and years. Following Williams, Miller & Ellner (2012), we checked for potential accidental eviction of individuals that would be projected to fall outside of the integration range [*L*, *U*]. Eviction of large adults occurred at values >*U*, and so we fixed U=180 mm to prevent accidental eviction. Extending the range of the upper integration limit did not qualititatively modify our results nor did it produce biologically unrealistic size transitions.

### Population simulations

To evaluate the threshold of population viability against projected climatic regimes, we examined the difference between the effect of current environmental conditions and simulated climate change conditions on the species’ population stochastic growth rate (*λ*_*s*_). Population trends were estimated by projecting the populations over a 100-year period in 100 separate runs with an initial population size *N* = 1,000 individuals. The initial population distribution of *N*, *n* in equation 1, was taken from the stable size distribution obtained from the kernel *K*. This is biologically meaningful because most sampled natural populations of this species are close to stationary equilibrium (*λ*=1, Rodríguez-Caro et al., 2019). Moreover, we carried out a sensitivity analysis to examine the role of the initial size distribution on our projections. However, comparisons of different initial size distributions did not significantly affect our estimates of stochastic growth rates under different climate scenarios (Appendix 3).

As we found that the best moving window model showed a significant effect of spring precipitation on the probability of reproduction *φ*, simulations of population viability included the effect of rainfall on *φ*. To simulate the effects of climate change on the viability of *T. graeca* populations, we quantified the current climate conditions, and then evaluated the effect of unusual increases in droughts on long-term demographic viability. The current regime of drought was estimated using Standardised Precipitation Index (SPI) from 1961 to 2018 (Vicente-Serrano et al., 2017). We defined the 10^th^ percentile of SPI (−1.2) as the limit of a drought. Thus, the current scenario is characterised by one drought per decade. The harsher scenarios of climate models predict increases in droughts due to a ~20% reduction of precipitations in SE Spain (Amblar, Casado-Calle, Pastor-Saavedra, Ramos-Calzado & Rodríguez-Camino, 2017), with a consequent 1.5 droughts per decade. Next, we projected an increase in the decadal recurrence of droughts to identify the threshold of the buffering ability of *T. graeca* against adverse conditions. We tested the current probability of drought from 1 to 7 droughts per decade. For the simulations, we categorised two states of precipitation for the study site for further analyses: (i) drought years at 61 mm of spring precipitation (10^th^ percentile of the historical records of precipitation, 1995 - 2017), and (ii) normal years at 104 mm of spring precipitation (computed as the average of the historical records of precipitation without the 10^th^ percentile).

For each of the projected simulation, we calculated the stochastic population growth rate (*λ*_*s*_) as:

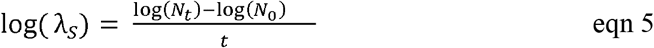

where *N_0_* is the population size at time *t* = 0 and *N_t_* the projected population size at time *t*. Population decreases when log(*λ*_*s*_) < 0 and increases when log(*λ*_*s*_) > 0, whereas log(*λ*_*s*_) = 0 indicates a demographically stable population (we evaluated the relation between droughts recurrence and log(*λ*_*s*_) using Pearson correlation).

We performed a perturbation analysis to identify the different contributions of the vital rates parameters to population growth rate *λ* (Griffith, 2017) under the different climate regimes: normal year and drought year, as defined above. We employed the brute-force method to calculate the elasticity of parameter-level vital rates to *λ* (Morris & Doak 2002). Briefly, this involves increasing the value of a particular vital rate parameter (*e.g*. intercept, slope) of interest by an infinitesimal amount (here 0.001) while keeping all other parameters unaltered and calculating the differential effect on *λ* compared to its original, unperturbed value. We chose to carry out this elasticity analysis at the vital rate parameter-level rather than higher levels like whole vital rate or IPM mesh-discretised bin-level because a vital rate parameter-level elasticity provides a greater insight into the underlying mechanics of size◻dependency in vital rates (Griffith, 2017). We estimated the confidence interval of the perturbation analysis using 100 bootstraps at individual-level resampling (Hall and Martin, 1988).

## Results

### Simulations of current scenario and the increase of drought recurrence

In our simulations of population performance under the current climatic regime, the population was demographically stable. Our model under the current climatic conditions of one drought per decade produced an estimate of stochastic population growth rate log(*λ*_*s*_) = 0.0000 (−0.0001 to 0.0000, 95% CI; Fig 2).

**Fig 2.**
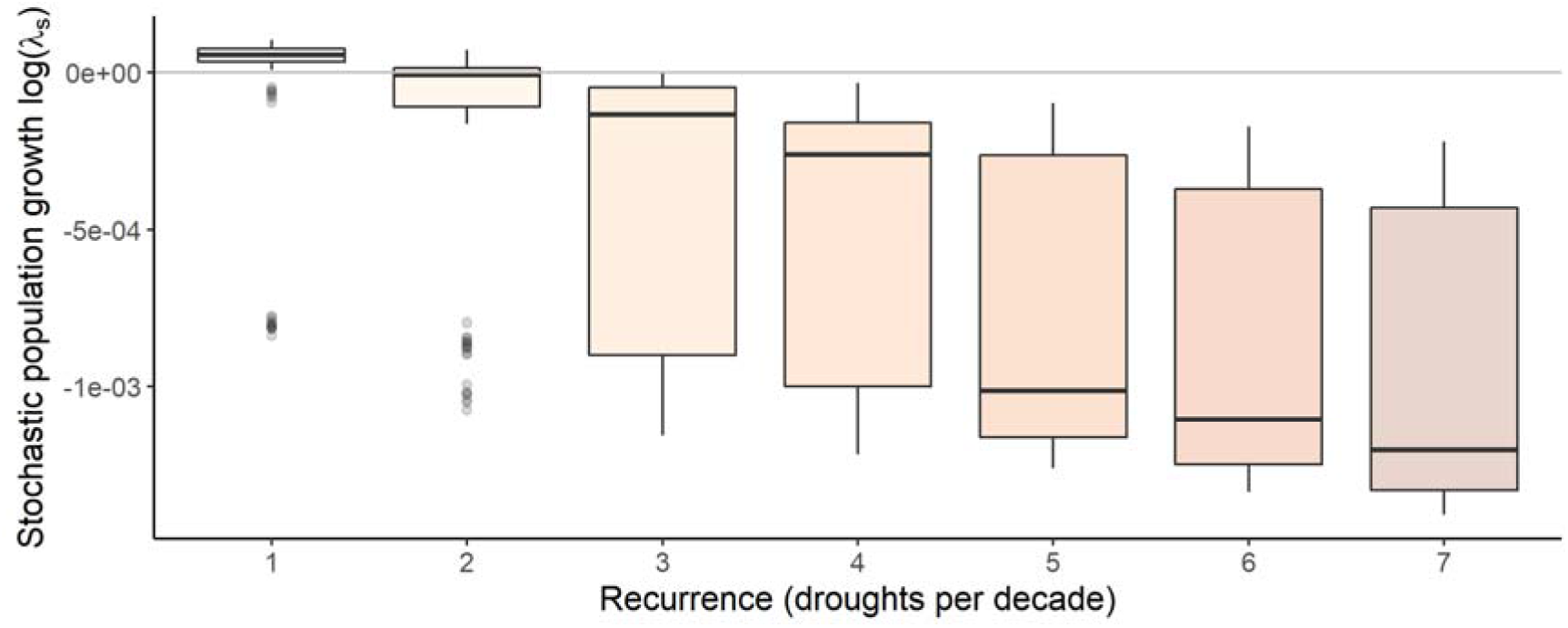
Stochastic population growth rates (log(*λ*_*s*_)) of the spur-thighed tortoise at the Galera population (Figure 1) under the current precipitation regimes (1 drought per decade) and under different drought recurrence intervals projected in the future. Boxes represent the interquartile range, the horizontal line represents the median, vertical line represents the upper and lower extreme values of the 95% of the interquartile range, and dots are the outlier values, defined as data outside this range. Grey line represents the stability of the populations (log(*λ*_*s*_) = 0). Any scenario where droughts occur three or more times per decade produce a statistically unviable stochastic population growth rate.

To explore the threshold of this tortoise species’ ability to demographically buffer against increasingly adverse conditions, we simulated different recurrence intervals of spring droughts per decade. The performance of the population under a projected scenario of an increase to two droughts per decade also resulted in demographic viability: log(*λ*_*s*_) = −0.0002 (−0.0003 to 0.0000, 95% CI). The simulated scenario characterised by a recurrence of three droughts per decade resulted in a negative stochastic population growth rate log(*λ*_*s*_) = −0.0004 (−0.0004 to −0.0003 CI), as was also the case for any scenario with a greater drought recurrence (Fig. 2). For instance, four and five droughts/decade produced values of log(*λ*_*s*_) of population decline, −0.0005 (−0.0006 to −0.0005, 95% CI) and −0.0007 (−0.0008 to −0.0006, 95% CI), respectively. A correlation between *λ*_*s*_ values and recurrence of droughts showed a significant negative relationship (Pearson correlation; *r* = 0.997, *p*<0.001; Fig. 2).

### Perturbation analyses

To understand the role of vital rates in the population growth rates of this endangered species under different climatic conditions, we examined the elasticity of the population growth rate *λ* at normal years (spring precipitation = 104 mm) and drought years (spring precipitation = 61 mm) to vital rates parameters. Under normal conditions, the slope of the growth (*γ slope* in Fig. 3) showed the largest and positive effect on *λ*, highlighting the importance to reach a reproductive size as soon as possible, instead of the initial size as a juvenile (*γ int* in Fig. 3). Next, the parameters with larger values of elasticity to *λ* were the quadratic slope of the size in the probability of reproduction (*φ slope 2*) and the slope of survival (*σ slope*). The relatively large elasticity of the quadratic slope of the probability of reproduction (*φ slope 2*) is a consequence of the variation in the range of female size with higher probability to reproduce. That is, small variation in this parameter highly modifies the shape of the curve of the probability of reproduction with an increment of the range of size values of females with higher probabilities of reproduction. On the other hand, the negative effect of the slope of the precipitation term on the probability of reproduction (*φ prec*) seems *a priori* counter-intuitive. However, this negative effect is due to the fact that the quadratic shape of the probability of reproduction describes the reproductive senescence in this species: high values in *φ prec* displace the optimal size for reproduction out of the boundaries of the observed size of these tortoises. The effects of the remaining parameters related to reproduction were negligible, as shown by the 95% C.I. bars in Fig. 3.

**Fig 3.**
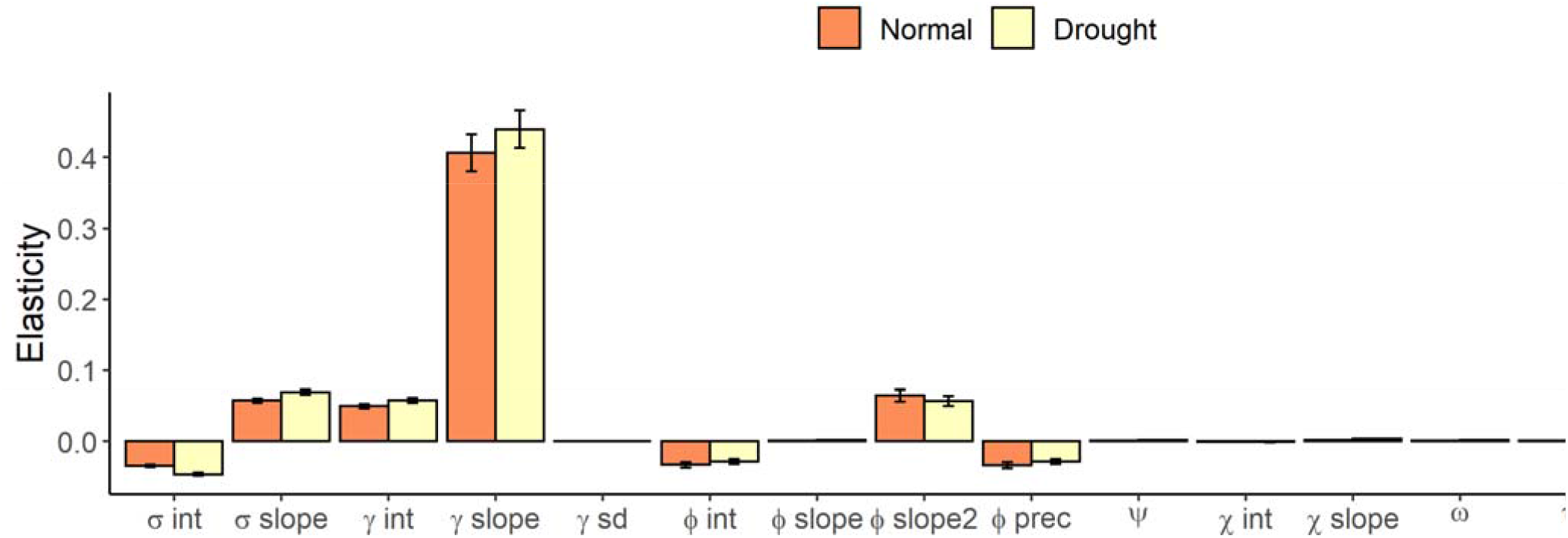
Elasticity of population growth rate *λ* with respect to the different vital rates parameters involved in the life cycle of *Testudo graeca* (Fig. 1b) under a regime of normal precipitation (orange) and drought (yellow). Normal years are defined as those with a spring precipitation of 104 mm, the average precipitation of the long-term weather data at the Galera population (Fig. 1a); Drought years are defined as those with a spring precipitation of 63 mm (10^th^ percentile at the same location). The vital rates parameters are: survival intercept (*σ* int), and slope (*σ* slope); growth intercept (*γ* int), slope (*γ* slope), and standard deviation (*γ* sd); probability of reproduction intercept (*φ* int), slope of linear size effect (*φ* slope), quadratic size effect (*φ* slope2) and precipitation effect (*φ* prec); the constant of number of clutches per year (*ψ*), number of eggs produced per adult per clutch intercept (*χ* int), and slope (*χ* slope); the probability of hatching (*ω*); the probability of surviving to the next year (*τ*); the mean size distribution of offspring of the next year (*θ* mean) and its standard deviation (*θ* sd).

The parameter elasticity profile for this species under a scenario of spring droughts remains largely similar to that under normal conditions (Fig. 3). However, key differences emerged in survival parameters under drought regime, where the survival intercept (*σ int*) became more negative and the survival slope (*σ slope*), which became even more positive when compared to the normal year (non-overlapping 95% CI). The emerging picture here, thus, is that during drought years survival becomes more important compared to its effect on *λ* during normal years. Although growth parameters increase the elasticity values, the growth rate intercept (*γ int*) showed higher elasticity under the drought regime than under a normal year (non-overlapping 95% CI), highlighting the importance of being larger as a juvenile under adverse conditions. Contrarily, the precipitation parameters related with the probability of reproduction decreased under drought regimes (*φ int*, *φ slope2* and *φ prec*). The remaining parameter elasticities showed small variations between drought and normal years.

## Discussion

Demographically buffered species cope with environmental stochasticity by allowing vital rates with low sensitivity to population growth rate to vary with environmental conditions (Pfister, 1998; Forcada, Trathan, & Murphy, 2008; McDonald et al., 2016), and deal with the challenges imposed by environmental variation (Morris et al., 2008; Villellas, Doak, García & Morris, 2015; Tavecchia et al. 2016). However, we hypothesised that future climate conditions, as predicted by climate change models (Hansen et al., 2006; Alexander et al., 2006; IPCC 2014), might render demographic buffering strategies ineffective beyond the threshold of variation. Our results support and identify the existence of a threshold beyond which our study species, the endangered *Testudo graeca*, cannot persist. In simulated scenarios when the environmental conditions were forced to reach thresholds of high recurrence of drought, three or more droughts per decade (defined as the 10^th^ percentile of historical records of precipitation), population dynamics showed negative, unviable trends.

The increase in extreme droughts frequency is one of the main threats of climate change for the biota (Burke et al., 2006; Seneviratne et al., 2012). Specially, drylands ecosystems in the Mediterranean Basin are highly threatened by climate change and desertification, with potentially irreversible impacts in (Maestre, Salguero-Gómez & Quero, 2012; Maestre, Soliveres, Gotelli, Quero & Berdugo, 2012). Such increases in the recurrence of adverse environmental conditions has been extensively documented to reduce the viability of populations (Beissinger, 1995), and to even cause their eventual extinction (Griffiths & Williams, 2000). Population collapses due to droughts have already been reported in several species, including amphibians (Walls, Barichivich & Brown 2013), mammals (Duncan, Chauvenet, McRae, & Pettorelli, 2012; Greenville, Wardle & Dickman, 2012), or invertebrates (Oliver et al., 2015). We also found that precipitation is the strongest driver of *T. graeca* reproduction and population can buffer the effects of drought, however our results suggest that the capability to buffer drought events has a limit. This limit became evident when the drought recurrence was greater than that predicted by harsher climate change scenarios.

As we expected for long lived species, growth and survival were key vital rates for the demographic performance of the species (Heppell, Caswell & Crowder, 2000), followed by parameters related to the probability of reproduction. The constant growth through the life of these animals was the most important demographic process to maintain the population growth rate. Indeed, growth patterns are central of life-history theory and influence life-history traits including survival and reproduction (Roff, 2002). Even in adverse conditions of drought, the growth parameter did not change the importance in the demographic performance of the species, since the size at maturity have been described as one of the most important rate for turtles and tortoise (Shine and Iverson, 1995). Meanwhile, survival rates showed lower effect than growth (unlike other chelonians; Cunnington & Brooks, 1996), but an increase in importance under drought conditions. In this drought scenario, variations in survival patterns have higher effect in *λ* than normal conditions, as a direct consequence of the buffering strategy of the species to maintain population growth rate (Heppel et al., 2000). Finally, the demographic processes related to reproduction with higher impact in the population growth rate were related to the probability of reproduction, *i.e.* the proportions of females that can reproduce. The drought reduced the probability of reproduction and also reduced the effect of the precipitation term in the variation of *λ*. The negative elasticity of the precipitation term in normal conditions is a consequence the change in the optimal size of reproduction. In that scenario, the probability of reproduction in the optimal size is lower and the effect of the precipitation term is lower too. Our results are an example about how trait-based demographic approach provides accurate insight about the life cycle of the species (Ozgul, Coulson, Reynolds, Cameron & Benton, 2012).

Despite the importance of existing theory to predict demographic responses to environmental variability, most of the existing studies have focused on the vital rates that most affect population growth rate, such as adult survival in the case of long-lived species (Rotella, Link, Chambert, Stauffer & Garrott, 2012). Few works have evaluated the effect of reductions in reproductive rates of long-lived species on the population viability (but see, Turner, Medica & Lyons, 1984, Jiménez-Franco et al., 2020). Thus, it is important to note that, in our simulations, environmental conditions only affect reproduction. However, negative effects of drought on survival (Longshore, Jaeger & Sappington, 2003) and population growth (Díaz-Paniagua, Andreu & Keller, 2001; Longshore et al., 2003; Mitro, 2003) have been described in other related species. Thus, our approach is likely conservative, since environmentally-driven reductions in adult survival in this species would likely heavily reduce population growth rates (Graciá et al., 2020).

Here, we have shown that the effect of climate change on the reproduction of the spur-thighed tortoise was driven via precipitation. Although our results support that the species is able to buffer the current levels of drought events, harsher environments or the interaction of other factors (such as high temperature or fire) could increase the risk of local extinction (Díaz-Paniagua, Andreu & Keller, 2006; Sanz-Aguilar et al., 2011). These harsher conditions could already be present in certain locations within the range of this species due microhabitat conditions, particularly in south-facing slopes (Quero, Villar, Marañón & Zamora, 2006, Zunzunegui et al., 2010) and could result in local extinctions. On the other hand, the synergic effects of droughts and fires have been described as a threat for endangered animal populations such us mammals (Hale et al., 2016) or reptiles (Lunney, Eby & O’Connell, 1991). The prediction in the South-Eastern Spain for climate change also includes an increase in the climate aridity, and this process will likely lead to more frequent wildfires (Mouillot, Rambal & Joffre, 2002). An increment in wildfires could affect the vulnerability of animal populations (Recher, Lunney and Matthews, 2009). However, according to Sanz-Aguilar et al. (2011), tortoises can coexist with fires at natural current fire frequencies (<1 fire every 20-30 years). Still, fire events are known to severely reduce survival and growth rates in spur-thighed tortoise populations (Rodríguez-Caro et al., 2013). Importantly, increases in temperature are well known to affect the sex ratio determination in the eggs of reptiles, with hotter years resulting in more females (Pieau, 1975). The sex-ratio bias in the offspring can significantly reduce the matting encounters for further generations, affecting its demographic viability (Graciá et al., 2020), or even the disappearance of one gender altogether (Janzen, 1994).

Our work contributes to a growing body of literature that uses demographic approaches on long◻term, detailed demographic monitoring, coupled with high-resolution climatic data and bibliographic information to investigate demographic responses to environmental variation (e.g. Gaillard et al., 1989; Campos et al., 2017; Paniw, Ozgul & Salguero◻Gómez, 2018). The challenges involved in measuring population dynamics from empirical studies of wildlife animals are numerous, often requiring various data inputs of different resources to obtain all the necessary parameters to adequately describe the life cycle of the species of interest (e.g. Sæther & Bakke, 2000, Forcada et al., 2008; Vélez-Espino et al., 2015; Graciá et al., 2020). The integration of multiple sources of detailed data into a single demographic model allowed us to look beyond the demographic effects of current environmental stochasticity and to reveal thresholds of demographic buffer capability of an endangered species.

Our modelling approach can be easily extrapolated to other long-lived demographically buffering species. To do so, researchers must carefully consider and ideally explicitly include means to evaluate and determine breakpoints for buffering such as interactive effects (e.g. temperature, disturbances; Darling & Côté, 2008) or temporal autocorrelation (Tuljapurkar & Haridas 2006; Paniw et al., 2018). Multiple environmental variables may interact to exacerbate -or cancel out in some cases (Paniw, Quintana◻Ascencio, Ojeda & Salguero◻Gómez, 2017)-their effects on natural populations (Darling & Côté, 2008). These variables can be explicitly included in the IPM simulations by testing their effects on each vital rate (e.g. Capdevila et al., 2019). On the other hand, temporal environmental autocorrelations, which realistically incorporate important aspects of future climatic projections (Paniw et al., 2018) can be included in these models using Markov chain approaches. In them, one identifies the different environmental states and the frequency of changes among them, couples them with demographic models parameterised under each environmental state and simulate different walks through these chains (Tuljapurkar & Haridas, 2006).

## Conclusions

This study contributes to our understanding of the ability of demographically buffering strategies to cope with stochastic environments and provides a useful modelling framework to predict population responses to environmental change. Moreover, through perturbation analyses, this framework allows us to identify the demographic mechanisms that mediate environmentally-driven population declines. Using this framework, our study provides empirical evidence about the limit for demographically buffering species. Although here we have found that *Testudo graeca* is currently demographically stable, a scenario of three or more extreme droughts per decade would jeopardise its long-term viability.

## Acknowledgements

The Spanish Ministry of Science and European Regional Development Fund funded this work through Project CGL2015-64144 (MINECIO/FEDER). P.C. was supported by a Ramón Areces Foundation Postdoctoral Scholarship (BEVP30P01A5816). JMB was supported by Juan de la Cierva contract (Ministerio de Economía y Competitividad, MEC; IJCI-2017-32149). R.S-G. was supported by a NERC IRF grant (NE/M018458/1). The Dirección General de Gestión del Medio Natural de la Junta de Andalucía (SGB/FOA/AFR) and the Dirección General de Medio Natural de la Comunidad Autónoma de la Región de Murcia have authorized and facilitated the follow-up work of the territories of the populations of *Testudo graeca* (SGB/FOA/AFR, SGYB/FOA/AFR/MVAC, SGYB/AF/DBP, SGYB/AFR/DBP, FA/AUT/CAP-06, AUT/ET/UND/48/2010, AUF20140057, AUF20160056).

