## Appendix 1 for "The demographic buffering strategy has a threshold of effectiveness to increases in environmental stochasticity"

**Appendix 1. Site monitoring**

**Table A1.** Number of females radiographed per site with their coordinates in UTM 30N across the seven years to estimate reproduction rates. Galera (in bold) was the site where survival and growth information were obtained (Sanz-Aguilar, 2011; Rodríguez-Caro et al., 2013), and so our climatic analyses and demographic simulations were carried out at that location. Due to limits imposed by the costs of monitoring, fieldwork did not take place in years 2008, 2009, 2012 and 2013**.**

| Site | X | Y | 2006 | 2007 | 2010 | 2011 | 2014 | 2015 | 2016 |
|---|---|---|---|---|---|---|---|---|---|
| **Galera** | **619500** | **4155500** | **7** | **41** | **59** | **35** | **43** | **15** | **38** |
| Adanes | 630500 | 4165500 | | | | | 43 | | |
| Alboluncas | 597500 | 4116500 | | | | | 12 | 8 | 14 |
| Bas Sur | 632500 | 4151500 | | | 10 | 29 | | | |
| Chinas | 597500 | 4119500 | | | 9 | 36 | | | |
| Chuecos | 624500 | 4154500 | | | 4 | 13 | | | |
| Culebras | 598500 | 4153500 | | | | | 21 | | 4 |
| Lomas del escribano | 622500 | 4150500 | | | | | 26 | | |
| Madroñales | 624500 | 4157500 | | | | 13 | | | |
| Marinica | 611500 | 4130500 | | | 7 | | | | |
| Palas | 608500 | 4131500 | 1 | | 5 | 5 | | | |
| Pisadas de la Virgen | 612500 | 4184500 | | | | | 9 | | |
| Sierrecica | 606500 | 4129500 | 6 | | 19 | 19 | | | |
| Tova | 603500 | 4180500 | | | | | | | 13 |
| Villaltas | 593500 | 4118500 | 9 | | 9 | 52 | | | |
