## Appendix 2 for "The demographic buffering strategy has a threshold of effectiveness to increases in environmental stochasticity"

**Appendix 2. Results of *climwin* analyses**

The results of the environmental analysis by *climwin* R package (Bailey & van de Pol, 2016) showed that models with precipitation as a covariate fitted the reproduction probability of the tortoise with a reduction of 62.69-52.32 the AICc of the null model (in the null model, the probability of reproduction is affected only by the size of tortoise individuals). The second variable with highest importance was temperature (45.58 – 31.90 AICc reduction) and NDVI (33.35 – 30.19 AICc reduction) (Table A2).

For precipitation, the temporal window that affected reproduction most prominently occurred in the spring (i.e., window starting three months before the reproduction event and finishing when the reproduction event had happened). We found a significant effect of precipitation during the previous summer (between 10 and 11 months before the reproduction). However, we did not use this variable because it is highly correlated with spring precipitation (Pearson correlation; $r = 0.823$, $p<0.001$).

**Table A2.** Model selection for the probability of reproduction with the three variables (precipitation, temperature and NDVI). *Window start* and *Window end* represent the period selected by the model, *ΔAICc* is the relation with the AICc of the null model (null model: probability of reproduction (*φ*) is affected just per individual size of the tortoise). *Effect* shows the effect of the time window on the variable (e.g. negative relation in the first model indicate that precipitation between august and September of the previous year is negative correlated with *φ*). For precipitation, we used accumulated rain; for temperature and NDVI, the average during the pertinent time window.

| Variable | Window start | Window end | ΔAICc | Effect |
|---|---|---|---|---|
| Precipitation | 11 | 10 | -62.69 | - |

| | | | | |
|---|---|---|---|---|
| | 2 | 1 | -62.57 | + |
| | 10 | 10 | -61.11 | - |
| | 3 | 1 | -61.08 | + |
| | 2 | 0 | -58.61 | + |
| | 2 | 2 | -65.36 | + |
| | 3 | 0 | -55.33 | + |
| | 3 | 2 | -52.32 | + |
| Temperature | 2 | 1 | -45.58 | - |
| | 3 | 1 | -45.41 | - |
| | 8 | 1 | -44.44 | - |
| | 12 | 1 | -41.89 | - |
| | 11 | 11 | -39.83 | + |
| | 2 | 0 | -38.86 | - |
| | 10 | 1 | -38.90 | - |
| | 11 | 1 | -37.90 | - |
| NDVI | 2 | 2 | -33.35 | + |
| | 8 | 2 | -30.95 | + |
| | 7 | 6 | -30.94 | + |
| | 8 | 6 | -30.82 | + |
| | 8 | 1 | -30.60 | + |
| | 2 | 1 | -30.51 | + |
| | 4 | 4 | -30.20 | + |
| | 7 | 2 | -30.19 | + |
