## Appendix 3 for "The demographic buffering strategy has a threshold of effectiveness to increases in environmental stochasticity"

**Appendix 3. Effect of initial size distribution on the population growth rates**

We carried out a sensitivity analyses to identify the effect of the initial size distribution on the stochastic population growth rate of the simulations. We assessed the initial size distribution for three different stable size distribution (SSD): SSD at demographic equilibrium ($SSD_{eq}$ used in the manuscript, which correspond to λ = 1.00), SSD for a negative growth rate ($SSD_{neg}$; λ = 0.98) and SSD for a positive growth rate ($SSD_{pos}$; λ = 1.02). The rationale of these choices is to maintain a real size distribution close to the stability as it was defined in previous studies (Rodríguez-Caro et al., 2019; Graciá et al., 2020), with a small range of variation of population growth rate (Δλ=0.02), as it was found in the natural populations (Rodríguez-Caro et al., 2016). We tested SSD from negative and positive population growth rate to address a high level of variation. We simulated them in seven scenarios of drought recurrence: one to seven droughts per decade. Then we tested if the stochastic growth ($\lambda_s$) was statistically significantly different across the initial SSD in using Kruskal Wallis test in R. Our results revealed similar outcomes for stochastic growth rates ($\log(\lambda_s)$) for three initial size distributions. We did not find differences in $\log(\lambda_s)$ among the initial $SSD_{eq}$, $SSD_{pos}$ or $SSD_{neg}$ in six scenarios of drought recurrence (two and seven droughts per decade; Table A3). We found statistical differences in the first scenario, one drought per decade (Table A3), but the values of $\log(\lambda_s)$ were equal, for the three initial SSD ($\log(\lambda_s)$ = 0.0000; -0.0001 to 0.0000, 95% CI), the differences appear in a small level, just appreciated in the remaining decimals.

**Table A3.** Results of Kruskal Wallis test among the stochastic growth lambda estimated with different initial stable size distribution (SSD) for each scenario. We compare the results of three initial SSD, in equilibrium ($SSD_{eq}$) and the initial stable size distribution for positive and negative population growth rates ($SSD_{pos}$ and $SSD_{neg}$, respectively). The analyses were carried out for seven scenarios of drought recurrence (one to seven droughts per decade). Results of the chi-square of Kruskal-Wallis tests ($\chi^2$), $p$ values for each test and degree of freedom (*d.f.*). In bold the significant results.

| Drought frequency per decade | $\chi^2$ | $p$ | *d.f.* |
|---|---|---|---|
| 1 | 14.96 | **<0.001** | 2 |
| 2 | 2.90 | 0.234 | 2 |
| 3 | 3.01 | 0.222 | 2 |
| 4 | 2.04 | 0.361 | 2 |
| 5 | 5.53 | 0.063 | 2 |
| 6 | 3.51 | 0.172 | 2 |
| 7 | 0.93 | 0.628 | 2 |
